# Abnormal trophoblast invasion in early onset preeclampsia: involvement of cystathionine β-synthase, specificity protein 1 and miR-22

**DOI:** 10.1101/2023.03.08.531728

**Authors:** Pallavi Arora, Sankat Mochan, Sunil Kumar Gupta, Neerja Rani, Pallavi Kshetrapal, Sadanand Dwivedi, Neerja Bhatla, Renu Dhingra

## Abstract

**Introduction:** Impaired trophoblast invasion has been observed in early onset Preeclampsia patients (EOPE). Trophoblast cell invasion during human placentation is majorly regulated by the balance between MMPs 2, 9 and their inhibitors [tissue inhibitors of matrix metalloproteinases (TIMPs 1, 2)]. Exogenous NaHS (hydrogen sulphide donor) treatment was shown to significantly increase the expression levels of matrix metalloproteinases (MMPs 2, 9) in human bladder cancer EJ cells. Epigentically, the gene expression of hydrogen sulphide synthesising enzyme CBS (cystathionine β-synthase) could be further regulated by various mi-RNAs via the transcription factors like Sp1. Specificity protein 1 (Sp1) has been identified as a target gene for miR-22 to regulate the invasion and metastasis of gastric cancer cells. However, the mechanism of MMPs regulation by either CBS, Sp1 and miRNA-22 in the pregnancies having EOPE is not known.

**Aims and Objectives:** To determine and compare the expression of MMPs 2, 9, TIMPs 1, 2, CBS, Sp1 and miRNA-22 in EOPE patients and normotensive, non-proteinuric controls.

**Materials and methods:** 100 pregnant women were enrolled from Department of Obstetrics and Gynaecology, AIIMS, New Delhi, India. EOPE women (n=50) after clinical diagnosis as per ACOG guidelines were enrolled as cases and normotensive, non-proteinuric pregnant women (n=50) were enrolled as controls. Protocol of the study was approved by Institute Ethics Committee, AIIMS, New Delhi. 5 ml of venous blood was collected from all the recruited women (2.5 ml in each EDTA and sera vial) followed by plasma and sera separation. Plasma samples were used subsequently to determine gene expression of MMPs 2, 9, TIMPs 1, 2, CBS, Sp1 and miRNA-22 by qRT-PCR and sera samples were used to estimate their protein levels by ELISA. Data were analyzed by STATA 14 and Graph Pad Prism 8.

**Results:** Significantly down regulated expression of MMPs 2, 9, CBS and Sp1 whereas up regulated expression for that of TIMPs 1, 2 was observed in EOPE patients as compared to healthy pregnant women at both transcription and translation levels. Expression of miR-22 (pre miR-22 and miR-22-3p) was found to be significantly elevated in EOPE patients as compared to normotensive, non-proteinuric controls.

**Conclusion:** This is the very first study of its kind which implicates that down regulated MMPs 2, 9, CBS, Sp1 levels and simultaneously upregulation of miR-22 expression in EOPE patients could have some association. In vitro experiments are needed to prove their association which if proven may provide a new, potential therapeutic target to treat early onset Preeclampsia.

## Introduction

Early-onset preeclampsia (EOPE) is a severe obstetrics disease which threatens mother and foetus^1^ and it develops within 34 weeks of pregnancy. Due to its early onset, EOPE has been the main cause of death of maternal women and perinatal infants. The pathogenesis of EOPE hasn’t been explained completely till date. In recent years, researches have shown that the failure of invasive ability of trophoblast cells is the crucial factor for pathogenesis of EOPE^2,3^. Placental extravillous trophoblasts rapidly proliferate, migrate and invade into myometrium and remodel maternal spiral arteries during implantation^4^. This process of placental invasion includes the expression of endopeptidases that degrade extracellular matrix (ECM) and help in tissue remodeling called matrix metalloproteinases (MMPs)^5^, especially MMP2 and MMP9, are known for their important role during pregnancy, being involved in the degradation of collagen IV, which is the prime component of maternal basement membrane^6^ and their expressions have been shown to mediate the invasiveness of cultured trophoblasts into the Matrigel^7^. Hence, MMP2 and MMP9 are regarded as the key enzymes during implantation. Tissue inhibitors of matrix metalloproteinases (TIMP1 and TIMP2) preferentially bind to both active and latent forms of MMP9 and MMP2^8^. The balanced expression of MMPs 2, 9 and TIMPs 1, 2 has profound implications for regulating the depth of trophoblast invasion into the uterus^9^. Exogenous NaHS (H_2_S donor) treatment significantly increased the expression levels of matrix metalloproteinases (MMPs 2, 9) in human bladder cancer EJ cells^10^. Endogenously, hydrogen sulphide (H_2_S) is produced as a metabolite of homocysteine (Hcy) by cystathionine β-synthase (CBS)^11^. It has already been documented that SL2 cells of drosophila, when transfected with mammalian Specificity protein 1 (Sp1) expression construct induced high levels of cystathionine β-synthase (CBS) activity indicating that Sp1 has a critical role in the regulation of CBS^12^. Sp1 was identified as a target gene for miR-22 to regulate the invasion and metastasis of gastric cancer cells^13^. miR-22 is a 22-nt noncoding RNA, was originally identified in HeLa cells as a tumor suppressing miRNA and was identified to be ubiquitously expressed in a variety of tissues^14^. The aim of the present study was to determine and compare the expressions of MMPs 2, 9, their inhibitors (TIMPs 2, 1), cystathionine β-synthase, specificity protein-1, precursor miR-22 and miR-22-3p in early onset preeclamptic (EOPE) patients and healthy pregnant women (normotensive, non-proteinuric).

## Materials and methods

### Study Subjects

100 pregnant women were enrolled from the antenatal clinic and the ward of the Department of Obstetrics and Gynaecology, All India Institute of Medical Science, New Delhi, India. Early onset preeclamptic women (n=50) after clinical diagnosis as per ACOG guidelines were enrolled as cases and normotensive, non-proteinuric pregnant women (n=50) (maternal and gestational age matched) were enrolled as controls. Protocol of the study was approved by the Institute Ethics Committee, AIIMS, New Delhi and written informed consent was obtained from all the enrolled women. 5 ml of venous blood was collected from all the recruited women (n=100) (2.5 ml in each EDTA and sera vial) followed by plasma and sera separation. Plasma samples were used subsequently to determine mRNA expression of MMPs 2, 9, TIMPs 1, 2, cystathionine beta synthase, specificity protein-1, precursor miR-22, miR-22-3p and sera samples were used to estimate the protein levels of MMPs 2, 9, TIMPs 1, 2, CBS and Sp1 in early onset preeclamptic patients and normotensive, non-proteinuric pregnant women.

### qRT-PCR (quantitative Real Time-Polymerase Chain Reaction)

RNA extraction was done from plasma by TRIzol reagent (Invitrogen). The quality of RNA was examined by denaturing gel and quantity was measured on Micro-Volume UV/Visible Spectrophotometer (Thermo Fisher Scientific-NanoDrop TM 2000). c-DNA preparation was done by Revert aid H-minus reverse transcriptase kit (Thermo). Quality of cDNA was checked on 0.8% agarose gel, visualized by ethidium bromide (EtBr) stain under UV and quantity was measured on Micro-Volume UV/Visible Spectrophotometer (Thermo Fisher Scientific-NanoDrop TM 2000). cDNA was amplified by quantitative RT-PCR (CFX96 Touch™ Real-Time PCR Detection System, BioRad). qRT-PCR reactions were carried out in 20 μl volume, including cDNA (template), SYBR Green (Thermo), forward and reverse primers (Sigma), and nuclease free water to determine mRNA expression of MMPs 2,9, TIMPs 1,2, CBS, Sp1 and gene expression of miR-22 (pre miR-22 and miR-22-3p). Glyceraldehyde-3-phosphate dehydrogenase (GAPDH) mRNA and U6 small nuclear RNA were used as internal controls. Primers were designed by NCBI and confirmed by In-Silico PCR (Table 1).

**Table 1:**
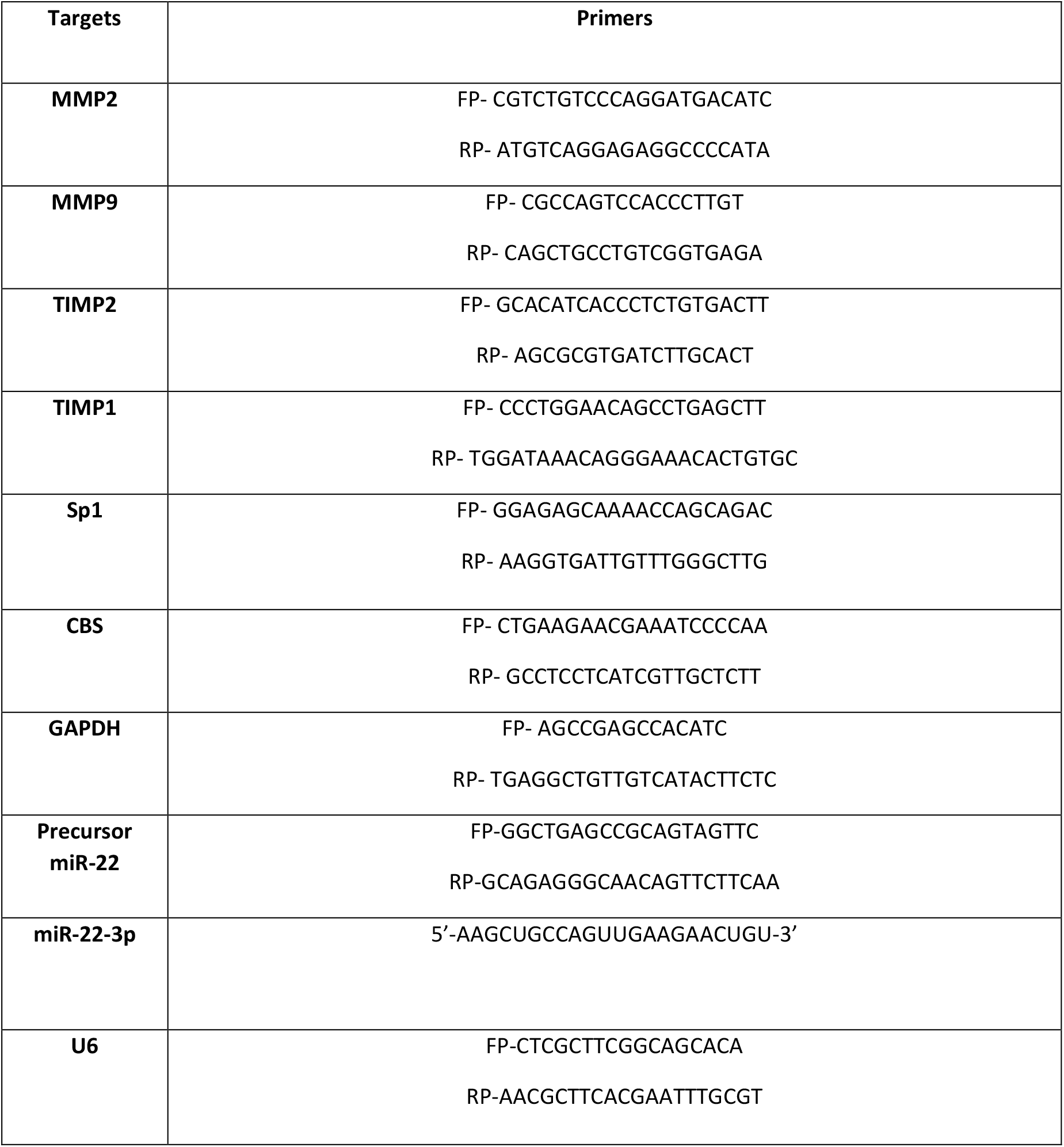
Primers designed by NCBI (National Centre for Biotechnology Information) and confirmed by In-Silico PCR. GAPDH: Glyceraldehyde 3-phosphate dehydrogenase

### ELISA

The levels of MMPs 2, 9, TIMPs 1, 2, CBS and Sp1 were estimated in the serum of early onset preeclamptic and normotensive pregnant women by sandwich ELISA [MMPs 2,9, TIMPs 1,2 (R & D Systems), CBS, Sp1 (G-Biosciences)].

### Statistical Analysis

Data was analyzed by STATA 14 and Graph Pad Prism 8. Relative quantification cycles of gene of interest (ΔCq) were calculated by ΔCq = Cq (target) - Cq (reference). Relative mRNA expression with respect to internal control gene was calculated by 2^-ΔCq^. Paired t-test and wilcoxon matched-pairs signed rank tests were used to compare the average level of the variable between two groups. *p* value<0.05 was considered statistically significant.

## Results

The clinical characteristics of early onset preeclamptic patients and normotensive, non-proteinuric controls are mentioned in Table 2.

**Table 2:** Clinical Characteristics of EOPE and normotensive, non proteinuric pregnant women (controls), n= number of subjects, Data presented as mean±SD, Paired t test, *statistical significance

| Study Groups |  |  |  |
| --- | --- | --- | --- |
| Clinical characteristics | EOPE (n=50) | Normotensive, Non proteinuric (Controls) (n=50) | Paired t test (p value) |
| Maternal age (years) | 29 ± 3.94 | 29.9 ± 4.44 | *p*>0.05 |
| Gestational age (weeks) | 30.68 ± 2.88 | 31.32 ± 3.65 | *p*>0.05 |
| Systolic blood pressure (mmHg) | 146.82 ± 10.59 | 115.26 ± 7.97 | *p*<0.0001* |
| Diastolic blood pressure (mmHg) | 95.8 ± 8.97 | 74.72 ± 7.51 | *p*<0.0001* |
| Protein (dipstick) | 27- 1+, 13- 2+, 10- 3+ | Nil or traces | NA |

### Significantly reduced expression of MMPs 2, 9 whereas elevated for those of TIMPs 1, 2 in EOPE patients

qRT-PCR and ELISA results showed that EOPE patients had significantly lower gene and protein expression of MMPs 2, 9 [Fig. 1(a,b), 2(a,b)]. On the other hand, TIMPs (1, 2) expression at both gene and protein levels found elevated in EOPE patients [Fig. 1 (c,d), 2(c,d)].

**Figure 1.**
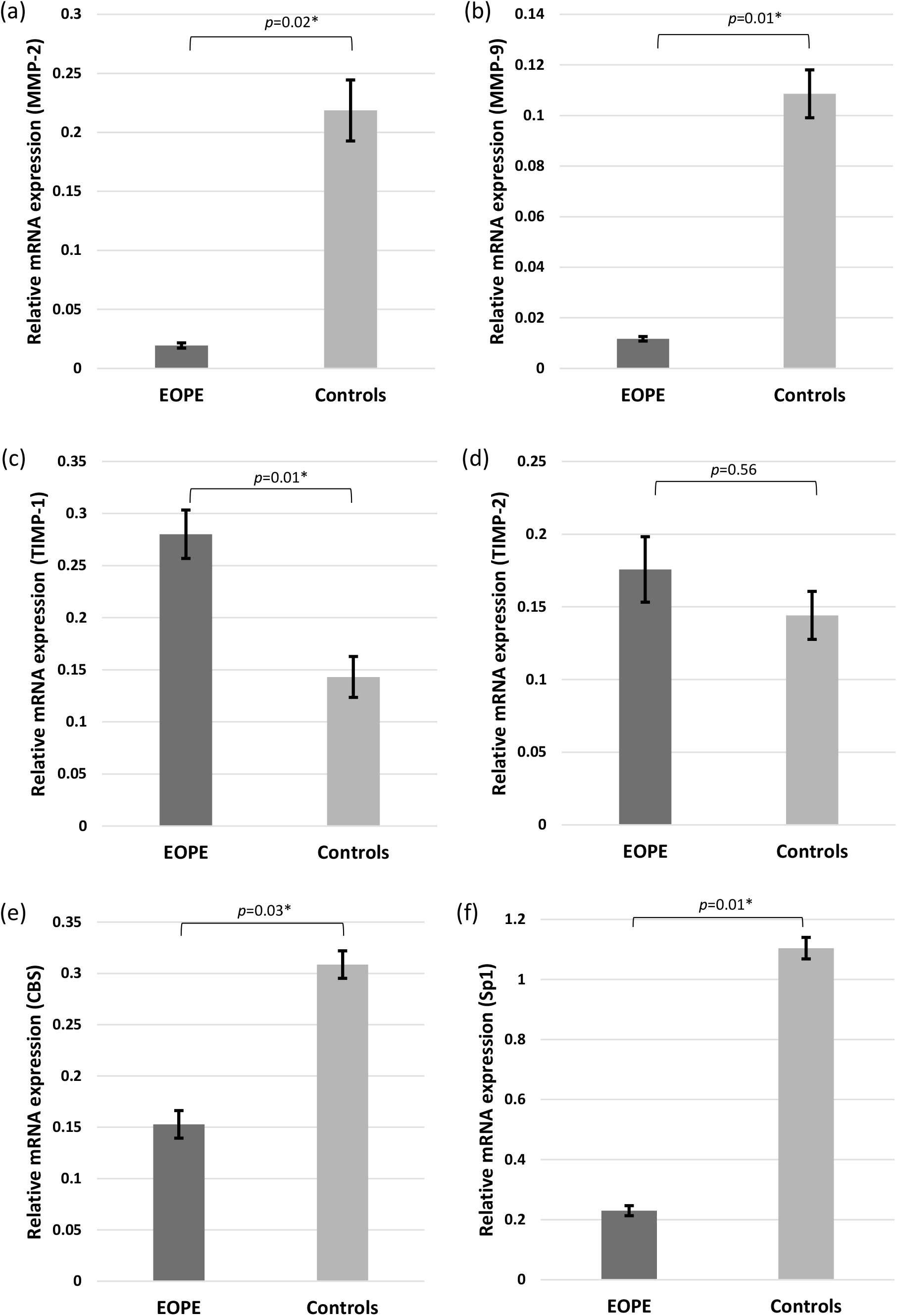
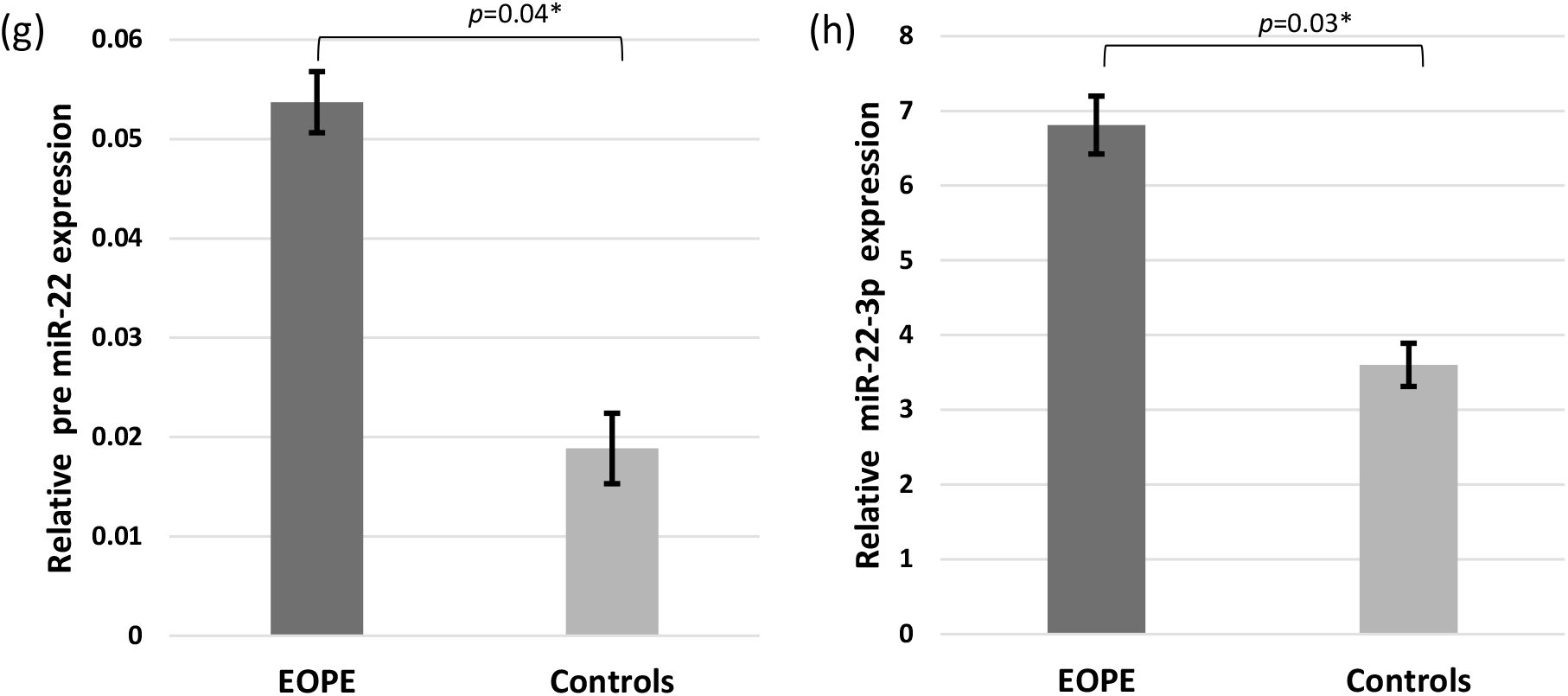
a-f: Bar diagrams represent the relative mRNA expression of MMPs 2, 9, TIMPs 1, 2, CBS and Sp1 in plasma samples of early onset preeclamptic patients and normotensive, non-proteinuric controls. GAPDH was used as positive control. g, h: Bar diagrams represent the relative precursor miR-22 and miR-22-3p expression. U6 was used as positive control. Data presented as mean ± SEM. Wilcoxon matched-pairs signed rank test was applied, *p* values indicated on graphs itself.

**Figure 2.**
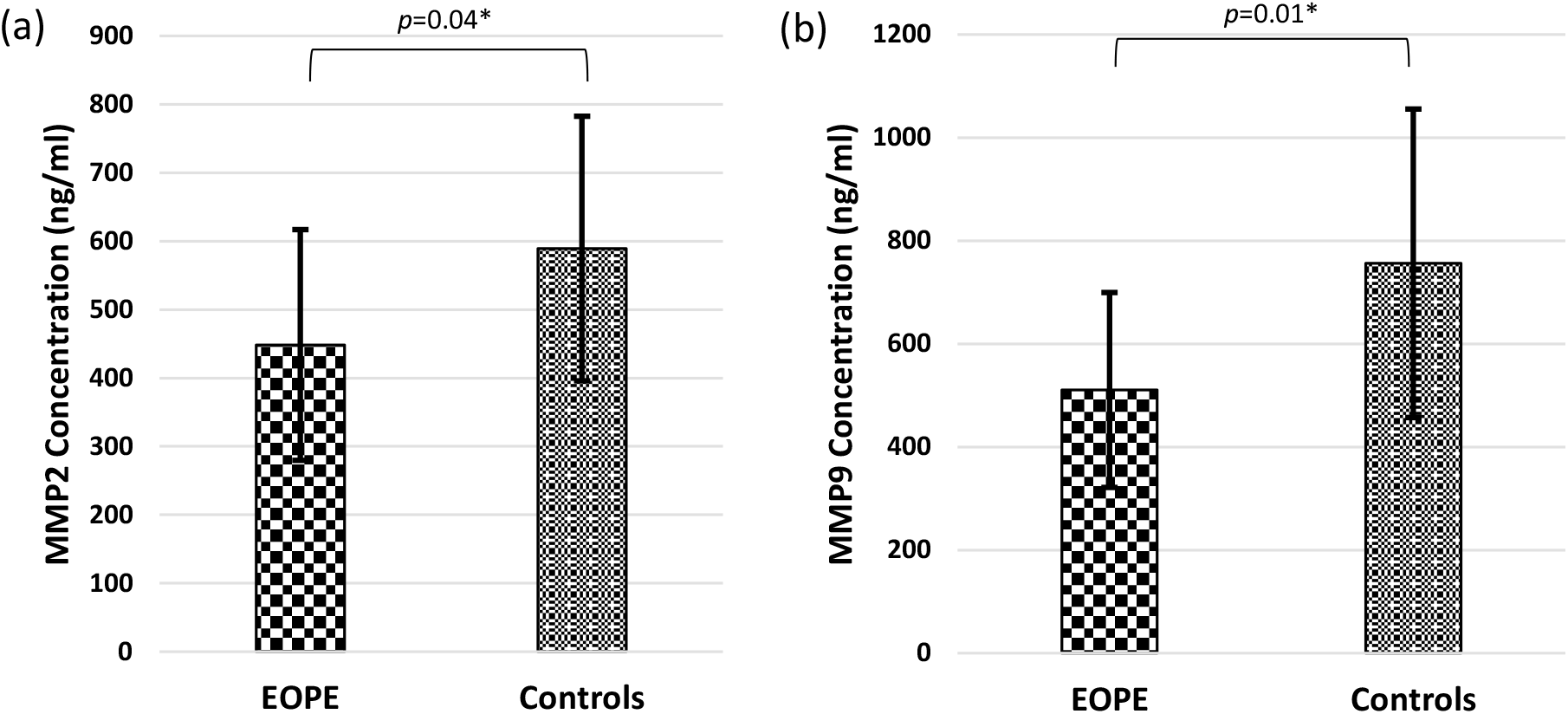
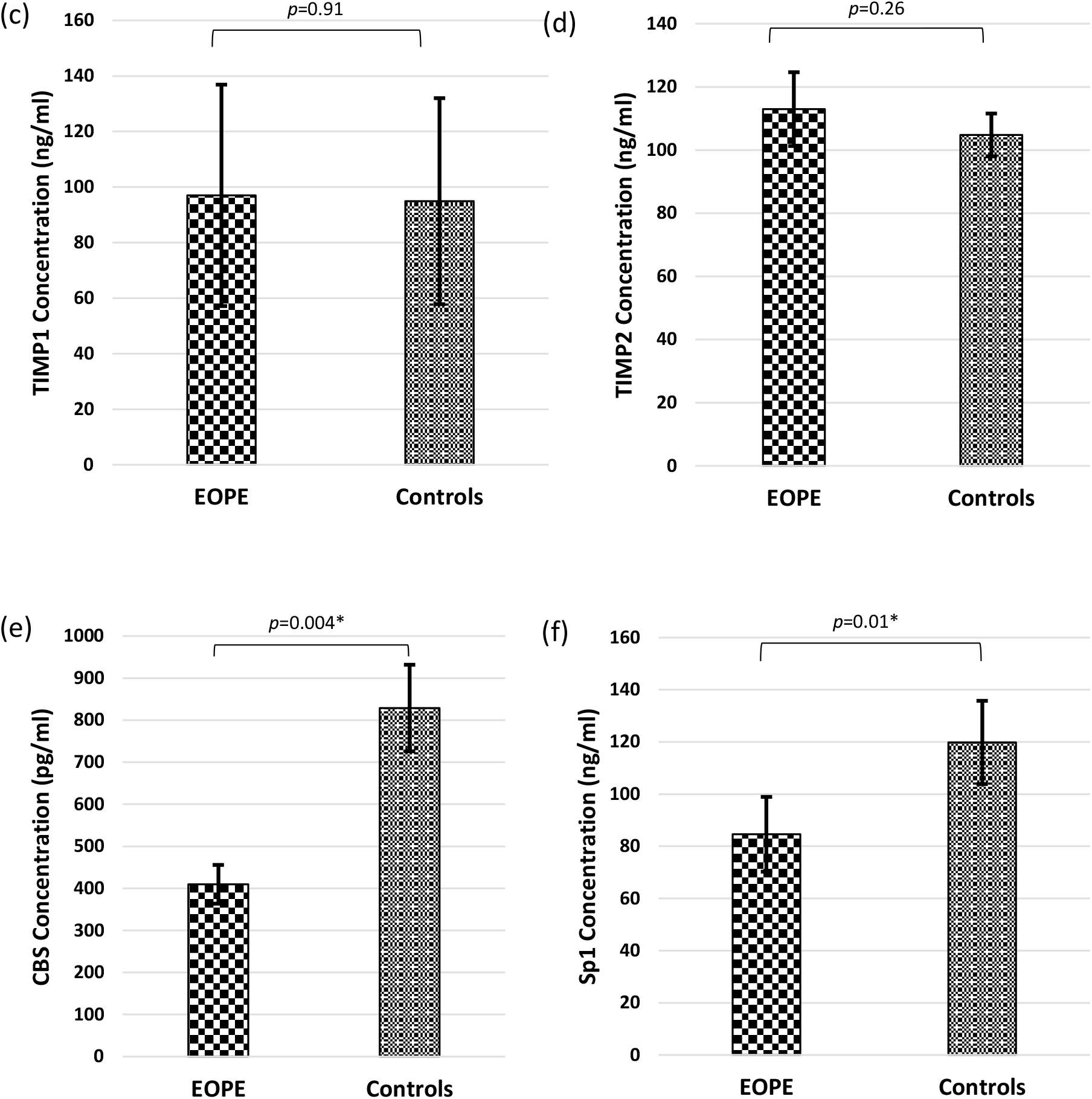
a-f: Bar diagrams represent serum levels of MMPs 2, 9, TIMPs 2,1, CBS and Sp1 proteins in sera samples of early onset preeclamptic patients and normotensive, non-proteinuric controls.. Data presented as mean ± SEM. Wilcoxon matched-pairs signed rank test was applied, *p* values indicated on graphs itself.

### Down regulated expression of Cystathionine β-synthase and Specificity protein 1 in EOPE patients

qRT-PCR and ELISA data unveiled that gene expression and protein levels of CBS [Fig. 1(e), 2(e)] and Sp1 [Fig. 1(f), 2(f)] were found to be significantly reduced in EOPE patients as compared to healthy pregnant women (normotensive non-proteinuric).

### Significantly elevated expression of precursor miR-22 and miR-22-3p in EOPE patients

qRT-PCR data revealed that patients with early onset preeclampsia showed significantly elevated plasma expression of miR-22 (pre miR-22 and miR-22-3p) than normotensive non-proteinuric pregnant women [Fig. 1 (g,h)].

## Discussion

Preeclampsia (PE) and eclampsia are estimated to cause over 50,000 maternal deaths worldwide per year^15^. In industrialized countries, rates of hypertensive disorders of pregnancy have risen, with African-American women at higher risk of associated mortality than American-Indian, white and Asian or Pacific-Islander women^16^. Women with PE or eclampsia have a 3–25 fold increased risk of severe complications in their index pregnancy, including abruption placentae, disseminated intravascular coagulation, pulmonary oedema and aspiration pneumonia^17^. Debate is ongoing regarding the heterogeneity of PE as the epidemiology, clinical presentation and associated morbidity differs between early onset or ‘placental’ preeclampsia (occurring before 34 weeks) and late onset or ‘maternal’ preeclampsia (occurring after 34 weeks)^18,19^. Early onset preeclampsia is associated with substantial risk of intrauterine growth restriction, whereas late onset disease is frequently associated with maternal obesity and large for gestational age neonates^20^. Abnormal placentation, reduced uterine perfusion pressure and placental ischemia are important initiating events in preeclampsia^21–23^. Inadequate placentation could be caused by abnormal inflammatory and immune responses and accumulation of natural killer (NK) cells and macrophages, trophosblast cells apoptosis, decreased invasion of spiral arteries, and abnormal expression of integrins and matrix metalloproteinases (MMPs) leading to decreased extracellular matrix (ECM) remodeling, shallow trophoblast invasion and poor spiral arteries remodeling^24^. MMP-2 and MMP-9 may be involved in ECM remodeling and trophoblast invasion of the spiral arteries during pregnancy^25–28^. The amount and activity of MMPs 2, 9 are increased in the aorta of normal pregnant rats, suggesting a role of MMPs in the pregnancy-associated vascular remodeling^29,30^. In the same line and length of the previous study, we also found significantly upregulated expression of MMPs 2, 9 in normal pregnant women as compared to early onset preeclamptic patients (EOPE) at both transcription and translation levels. On the other hand, MMPs expression/activity could be influenced by tissue inhibitors of metalloproteinases (TIMPs)^31–34^, for instance, TIMP-2 or specific MMP-2 blocking antibody inhibits cytotrophoblast invasion^28^. TIMP-1 is a tissue inhibitory factor corresponding to MMP-9 and the balanced expression of MMP-9 and TIMP-1 has extensive implications for regulating the depth of trophoblast invasion into the uterus^9^. In the present study, TIMPs 1, 2 expression was found upregulated in EOPE patients at both mRNA and protein levels which could have possibly resulted in defective cytotrophoblastic invasion leading to abnormal spiral artery remodelling in such patients.

Liu *et al*. in 2017 observed that exogenous H_2_S may promote cell proliferation and invasion by up regulating the expression level of MMP-2 and MMP-9 in human bladder cancer EJ cells^10^. In their study, different concentrations of exogenous NaHS (H_2_S donor) were applied to treat human bladder cancer EJ cells and the results showed that exogenous NaHS could promote both cell proliferation and invasion abilities, both of which were increased synchronously with the increased NaHS concentrations^10^. H_2_S is the gas signaling molecule observed in human body, which is generated by essential amino-acid cysteine in vivo through one carbon unit metabolism and transfer-sulfur pathway with enzyme catalysis and the included enzymes were mainly cystathionine β-synthase (CBS), cystathionine γ-lyase (CTH), and 3-mercaptopyruvate sulfurtransferase (MPST)^35–38^. We, in our study have observed significantly up regulated expression of CBS likewise MMPs 2, 9 in healthy pregnant women as compared to EOPE patients at both transcription and translation levels. To understand the molecular mechanism, there is a previous study by Kraus *et al*. in which they have determined the complete genomic sequence of human CBS^39^. Another study by the same group mapped the transcriptional start sites of five human CBS mRNA isoforms, designated CBS -1a, -1b, -1c, -1d, and -1e.^40^. Maclean *et al*. in 2004 found that isoforms -1a and -1b form the vast majority of transcripts, whereas isoforms -1c, -1d, and -1e are relatively rare^12^. They observed that both CBS -1a and -1b promoters are transactivated by specificity protein 1 (Sp1) and specificity protein 3 (Sp3) in Drosophila SL2 cells^12^. They also speculated that Sp1 is necessary and sufficient for growth specific regulation of CBS -1b promoter and the possible mechanism which they have explained was Sp1 and Sp3 form a ternary complex with each other prior to binding the CBS -1b promoter region and Sp1 induces conformational changes in CBS-1b promoter region that facilitate collateral binding of Sp3 with a resultant increase in CBS promoter activity^12^. We, in the present study observed significantly up regulated expression of Specificity protein 1 in healthy pregnant women as compared to EOPE patients at both transcription and translation levels. The elevated Sp1 expression could have led to the up regulation of CBS and MMPs at both gene and protein levels in healthy pregnant women as compared to EOPE patients.

Sp1 is identified as a transcriptional activator for various genes involved in almost all cellular processes in mammalian cells^41^, takes part in cancer development and progression^42,43^. Guo *et al*. in 2013 found that Sp1 is a putative target gene for miR-22, mediating cell migration and invasion^13^. They observed an inverse correlation between gene expression of miR-22 and Sp1 in the gastric cancer tissues^13^. In the present study, we observed up regulated miR-22 (pre miR-22and miR-22-3p) and down regulated mRNA expression of Sp1 in EOPE patients as compared to normotensive, non-proteinuric controls. Our findings are in line and length with the previous study.

## Summary

In conclusion, the down-regulation of MMPs 2, 9, CBS and Sp1 and simultaneous upregulation of TIMPs 1, 2 and miR-22 (pre miR-22 and miR-22-3p) expression was observed in patients which could have led to the development of EOPE. Further understanding of the interaction between MMPs 2, 9, CBS, Sp1 and miRNA should help design more specific and efficient measures for early detection of early onset preeclampsia.

## References

1. Vaiman, D., Calicchio, R. & Miralles, F. Landscape of transcriptional deregulations in the preeclamptic placenta. PLoS One., 8, pp. e65498 (2013).

2. Mayhew, T.M. Estimating oxygen diffusive conductances of gas-exchange systems: A stereological approach illustrated with the human placenta. Ann Anat., 196, pp. 34–40 (2014).

3. Basak, S. & Duttaroy, A.K. Effects of fatty acids on angiogenic activity in the placental extravillious trophoblast cells. Prostaglandins Leukot Essent Fatty Acids., 88, pp. 155–162 (2013).

4. Pijnenborg, R., Robertson, W.B., Brosens, I. & Dixon, G. Review article: trophoblast invasion and the establishment of haemochorial placentation in man and laboratory animals. Placenta., 2, pp. 71–91 (1981).

5. Soundararajan, R. & Rao, A.J. Trophoblast ‘pseudo-tumorigenesis’: significance and contributory factors. Reproductive Biology and Endocrinology., 2:15 (2004).

6. Staun-Ram, E. & Shalev, E. Human trophoblast function during the implantation process. Reproductive Biology and Endocrinology., 3(56) (2005).

7. Fisher, S.J., Cui, T.Y., Zhang, L., Hartman, L., Grahl, K., Zhang, G.Y., Tarpey, J. & Damsky, C.H. Adhesive and degradative properties of human placental cytotrophoblast cells in vitro. Journal of Cell Biology., 109, pp. 891–902 (1989).

8. Nagase, H., Visse, R. & Murphy, G. Structure and function of matrix metalloproteinases and TIMPs. Cardiovascular Research., 69, pp. 562–573 (2006).

9. Zhang, Y., Li, P., Guo, Y., Liu, X. & Zhang, Y. MMP-9 and TIMP-1 in placenta of hypertensive disorder complicating pregnancy. Experimental and therapeutic medicine., 18, pp. 637–641 (2019).

10. Liu, H., Chang, J., Zhao, Z., Li, Y. & Hou, J. Effects of exogenous hydrogen sulfide on the proliferation and invasion of human bladder cancer cells. J Can Res Ther.,13, pp. 829–32 (2017).

11. Sen, U., Sathnur, P.B., Kundu, S., Givvimani, S., Coley, D.M., Mishra, P.K., Qipshidze, N., Tyagi, N., Metreveli, N. & Tyagi, S.C. Increased endogenous H2S generation by CBS, CSE, and 3MST gene therapy improves ex vivo renovascular relaxation in hyperhomocysteinemia. Am J Physiol Cell Physiol., 303, pp. C41–C51 (2012).

12. Maclean, K.N., Kraus, E. & Kraus, J.P. The dominant role of Sp1 in regulating the cystathionine beta-synthase-1a and -1b promoters facilitates potential tissue-specific regulation by Kruppel-like Factors. The Journal of Biological Chemistry., 279(10), pp. 8558–8566 (2003).

13. Guo, M.M., Hu, L.H., Wang, Y.Q., Chen, P., Huang, J.G., Lu, N., He, J.H. & Liao, C.G. miR-22 is down-regulated in gastric cancer, and its overexpression inhibits cell migration and invasion via targeting transcription factor Sp1. Med Oncol., 30, pp. 542 (2013).

14. Xiong, J., Du, Q. & Liang, Z. Tumor-suppressive microRNA-22 inhibits the transcription of E-box-containing c-Myc target genes by silencing c-Myc binding protein. Oncogene.,29, pp. 4980–8 (2010).

15. Ghulmiyyah, L. & Sibai, B. Maternal mortality from preeclampsia/eclampsia. Semin. Perinatol. 36, 56–59 (2012).

16. Shahul, S., Tung, A., Minhaj, M., Nizamuddin, J., Wenger, J., Mahmood, E., Mueller, A., Shaefi, S., Scavone, B., Kociol, R.D., Talmor, D. & Rana, S. Racial disparities in comorbidities, complications, and maternal and fetal outcomes in women with preeclampsia/eclampsia. Hypertens. Pregnancy., 34, pp. 506–515 (2015).

17. Zhang, J., Meikle, S. & Trumble, A. Severe maternal morbidity associated with hypertensive disorders in pregnancy in the United States. Hypertens. Pregnancy., 22, pp. 203–212 (2003).

18. Robillard, P. Y., Dekker, G., Iacobelli, S. & Chaouat, G. An essay of reflection: why does preeclampsia exist in humans, and why are there such huge geographical differences in epidemiology? J. Reprod. Immunol., 114, pp. 44–47 (2016).

19. Lisonkova, S. & Joseph, K. S. Incidence of preeclampsia: risk factors and outcomes associated with early-versus late-onset disease. Am. J. Obstet. Gynecol.,209, pp. 544.e1–544.e12 (2013).

20. Rasmussen, S., Irgens, L. M. & Espinoza, J. Maternal obesity and excess of fetal growth in pre-eclampsia. BJOG., 121, pp. 1351–1357 (2014).

21. Khalil, R.A. & Granger, J.P. Vascular mechanisms of increased arterial pressure in preeclampsia: lessons from animal models. American journal of physiology. Regulatory, integrative and comparative physiology., 283(1), pp. R29–45 (2002).

22. Gilbert, J.S., Babcock, S.A. & Granger, J.P. Hypertension produced by reduced uterine perfusion in pregnant rats is associated with increased soluble fms-like tyrosine kinase-1 expression. Hypertension., 50(6), pp. 1142–1147 (2007).

23. Alexander, B.T., Kassab, S.E., Miller, M.T., Abram, S.R., Reckelhoff, J.F., Bennett, W.A. & Granger JP. Reduced uterine perfusion pressure during pregnancy in the rat is associated with increases in arterial pressure and changes in renal nitric oxide. Hypertension., 37(4), pp. 1191–1195 (2001).

24. Chen, J. & Khalil, R.A. Matrix Metalloproteinases in Normal Pregnancy and Preeclampsia. Prog Mol Biol Transl Sci., 148, pp. 87–165 (2017);

25. Shimonovitz, S., Hurwitz, A., Dushnik, M., Anteby, E., Geva-Eldar, T. & Yagel, S. Developmental regulation of the expression of 72 and 92 kd type IV collagenases in human trophoblasts: a possible mechanism for control of trophoblast invasion. American journal of obstetrics and gynecology., 171(3), pp. 832–838 (1994).

26. Suman, P. & Gupta SK. Comparative analysis of the invasion-associated genes expression pattern in first trimester trophoblastic (HTR-8/SVneo) and JEG-3 choriocarcinoma cells. Placenta., 33(10), pp. 874–877 (2012).

27. Su, M.T., Tsai, P.Y., Tsai, H.L., Chen, Y.C. & Kuo, P.L. miR-346 and miR-582-3p-regulated EG-VEGF expression and trophoblast invasion via matrix metalloproteinases 2 and 9. Biofactors., 43(2), pp. 210–219 (2017).

28. Isaka, K., Usuda, S., Ito, H., Sagawa, Y., Nakamura, H., Nishi, H., Suzuki, Y., Li, Y.F. & Takayama, M. Expression and activity of matrix metalloproteinase 2 and 9 in human trophoblasts. Placenta., 24(1), pp. 53–64 (2003).

29. Dang, Y., Li, W., Tran, V. and Khalil, R.A. EMMPRIN-mediated induction of uterine and vascular matrix metalloproteinases during pregnancy and in response to estrogen and progesterone. Biochemical pharmacology., 86(6), pp. 734–747 (2013).

30. Yin, Z., Sada, A.A., Reslan, O.M., Narula, N. and Khalil, R.A. Increased MMPs expression and decreased contraction in the rat myometrium during pregnancy and in response to prolonged stretch and sex hormones. Am J Physiol Endocrinol Metab., 303(1), pp. E55–70 (2012).

31. Raffetto, J.D. & Khalil, R.A. Matrix metalloproteinases and their inhibitors in vascular remodeling and vascular disease. Biochem Pharmacol., 75(2), pp. 346–359 (2008).

32. Visse, R. & Nagase, H. Matrix metalloproteinases and tissue inhibitors of metalloproteinases: structure, function, and biochemistry. Circ Res., 92(8), pp. 827–839 (2003).

33. Kucukguven, A. & Khalil, R.A. Matrix metalloproteinases as potential targets in the venous dilation associated with varicose veins. Curr Drug Targets., 14(3), pp. 287–324. (2013).

34. Pascual, G., Rodriguez, M., Gomez-Gil, V., Trejo, C., Bujan, J. & Bellon, J.M. Active matrix metalloproteinase-2 upregulation in the abdominal skin of patients with direct inguinal hernia. Eur J Clin Invest. 40(12), pp. 1113–1121 (2010).

35. Kimura, H. Hydrogen sulfide: Its production, release and functions. Amino Acids., 41, pp. 113–21 (2011).

36. Pandey, S. Hydrogen sulfide: A new node in the abscisic acid-dependent guard cell signaling network? Plant Physiol., 166, pp. 1680–1681 (2014).

37. Hancock, J.T. & Whiteman, M. Hydrogen sulfide and cell signaling: Team player or referee? Plant Physiol Biochem., 78, pp. 37–42 (2014).

38. Li, L., Rose, P. & Moore, P.K. Hydrogen sulfide and cell signaling. Annu Rev Pharmacol Toxicol., 51, pp. 169–187 (2011).

39. Kraus, J. P., Oliveriusova, J., Sokolova, J., Kraus, E., Vlcek, C., de Franchis, R., Maclean, K. N., Bao, L., Bukovska, G., Patterson, D., Paces, V., Ansorge, W. & Kozich, V. Genomics., 52, pp. 312–324 (1998).

40. Bao, L., Vlcek, C., Paces, V. & Kraus, J. P. Arch. Biochem. Biophys., 350, pp. 95–103 (1998).

41. Cawley, S., Bekiranov, S., Ng, H.H., Kapranov, P., Sekinger, E.A., Kampa, D. et al. Unbiased mapping of transcription factor binding sites along human chromosomes 21 and 22 points to widespread regulation of noncoding RNAs. Cell., 116, pp. 499–509 (2004).

42. Yao, J.C., Wang, L., Wei, D., Gong, W., Hassan, M., Wu, T.T. et al. Association between expression of transcription factor Sp1 and increased vascular endothelial growth factor expression, advanced stage, and poor survival in patients with resected gastric cancer. Clin Cancer Res., 10, pp. 4109–17 (2004).

43. Yuan, P., Wang, L., Wei, D., Zhang, J., Jia, Z., Li, Q. et al. Therapeutic inhibition of Sp1 expression in growing tumors by mithramycin a correlates directly with potent antiangiogenic effects on human pancreatic cancer. Cancer., 110, 2682–2690 (2007).

