## Supplementary file for "Abnormal trophoblast invasion in early onset preeclampsia: involvement of cystathionine β-synthase, specificity protein 1 and miR-22"

**ACOG Guidelines**

Blood Pressure- 140 mm Hg systolic or >90mm Hg diastolic on 2 occasions at least 4 hours apart after 20 weeks of gestational age in women with a previously normal BP and 160 mm Hg systolic or >110 mm Hg diastolic, confirmed within a short interval (minutes) to facilitate timely antihypertensive therapy; Proteinuria >300 mg per 24-hour urine collection or protein/ creatinine ratio > 0.3mg/dl or dipstick reading of >1+ or in the absence of proteinuria, new-onset hypertension with new onset of one or more of the following- thrombocytopenia: platelet count <100,000/µl, renal insufficiency: serum creatinine > 1.1mg/dl or doubling of serum creatinine in the absence of other renal disease, impaired liver function: elevated blood levels of liver transaminases to twice normal concentrations, pulmonary edema and cerebral edema.
